# From circuits to lifespan: translating mouse and human timelines with neuroimaging based tractography

**DOI:** 10.1101/2024.07.28.605528

**Authors:** Nicholas C. Cottam, Kwadwo Ofori, Madison Bryant, Jessica R. Rogge, Khan Hekmatyar, Jianli Sun, Christine J. Charvet

## Abstract

Age is a major predictor of developmental processes and disease risk, but humans and model systems (e.g., mice) differ substantially in the pace of development and aging. The timeline of human developmental circuits is well known. It is unclear how such timelines compare to those in mice. We lack age alignments across the lifespan of mice and humans. Here, we build upon our Translating Time resource, which is a tool that equates corresponding ages during development. We collected 477 time points (n=1,132 observations) from age-related changes in body, bone, dental, and brain processes to equate corresponding ages across humans and mice. We acquired high-resolution diffusion MR scans of mouse brains (n=12) at sequential stages of postnatal development (postnatal day 3, 4, 12, 21, 60) to trace the timeline of brain circuit maturation (e.g., olfactory association pathway, corpus callosum). We found heterogeneity in white matter pathway growth. The corpus callosum largely ceases to grow days after birth while the olfactory association pathway grows through P60. We found that a P3 mouse equates to a human at roughly GW24, and a P60 mouse equates to a human in teenage years. Therefore, white matter pathway maturation is extended in mice as it is in humans, but there are species-specific adaptations. For example, olfactory-related wiring is protracted in mice, which is linked to their reliance on olfaction. Our findings underscore the importance of translational tools to map common and species-specific biological processes from model systems to humans.

**Significance statement:** Mice are essential models of human brain development, but we currently lack precise age alignments across their lifespan. Here, we equate corresponding ages across mice and humans. We utilize high-resolution diffusion mouse brain scans to track the growth of brain white matter pathways, and we use our cross-species age alignments to map the timeline of these growth patterns from mouse to humans. In mice, olfactory association pathway growth continues well into the equivalent of human teenage years. The protracted development of olfactory association pathways in mice aligns with their specialized sense of smell. The generation of translational tools bridges the gap between animal models and human biology while enhancing our understanding of developmental processes generating variation across species.

## Introduction

The brain consists of a complex network of fiber bundles connecting different regions. In humans, some brain pathways continue to mature and rewire well into adulthood (Arain et al., 2013), while other pathways mature relatively earlier a few years after birth (Cohen et al., 2016; Mohammad et al., 2017). Fiber bundles that grow for an extended time are often associated with higher-level cognition, and their development is concomitant with behavioral changes during adolescence (Arain et al., 2013; Qu et al., 2015; Cohen et al., 2016). It is well-established that human development, including brain growth, proceeds more slowly than it does in many other mammals (Clancy et al., 2001; Workman et al., 2013; Charvet et al., 2023). Whether the timeline of fiber bundle growth is unusually protracted in humans relative to other species remains uncertain. We have yet to map the timetable of white matter pathway maturation in mice and compare these timelines to those found in humans. Such a comparison would necessitate age alignments across postnatal ages in humans and mice.

Here, we build on a long-standing project called Translating Time, which collects time points (e.g., when different neurons are generated, when eyes open) to identify corresponding ages across humans and various model systems (Figures 1-2, Clancy et al., 2001, 2007; Workman et al., 2013; Charvet, 2021). Previously, we developed an integrative approach to generate cross-species age alignments across the lifespan. We used several metrics to collect time points across several scales (e.g., anatomy, behavior, gene expression), to find age alignments across the lifespan of humans and great apes (Charvet et al., 2023). Here, we expand on this approach to collect time points. We use a range of metrics to find corresponding ages across humans and mice. Corresponding time points are extracted from bone, brain, and tooth maturation, and from temporal changes in gene expression. In humans, we collected time points up to their 50s and 60s. In mice, we collected time points into their 3rd year. We used these cross-species age alignments to compare the timeline of maturation in mouse brain circuit maturation to that of humans.

**Figure 1.**
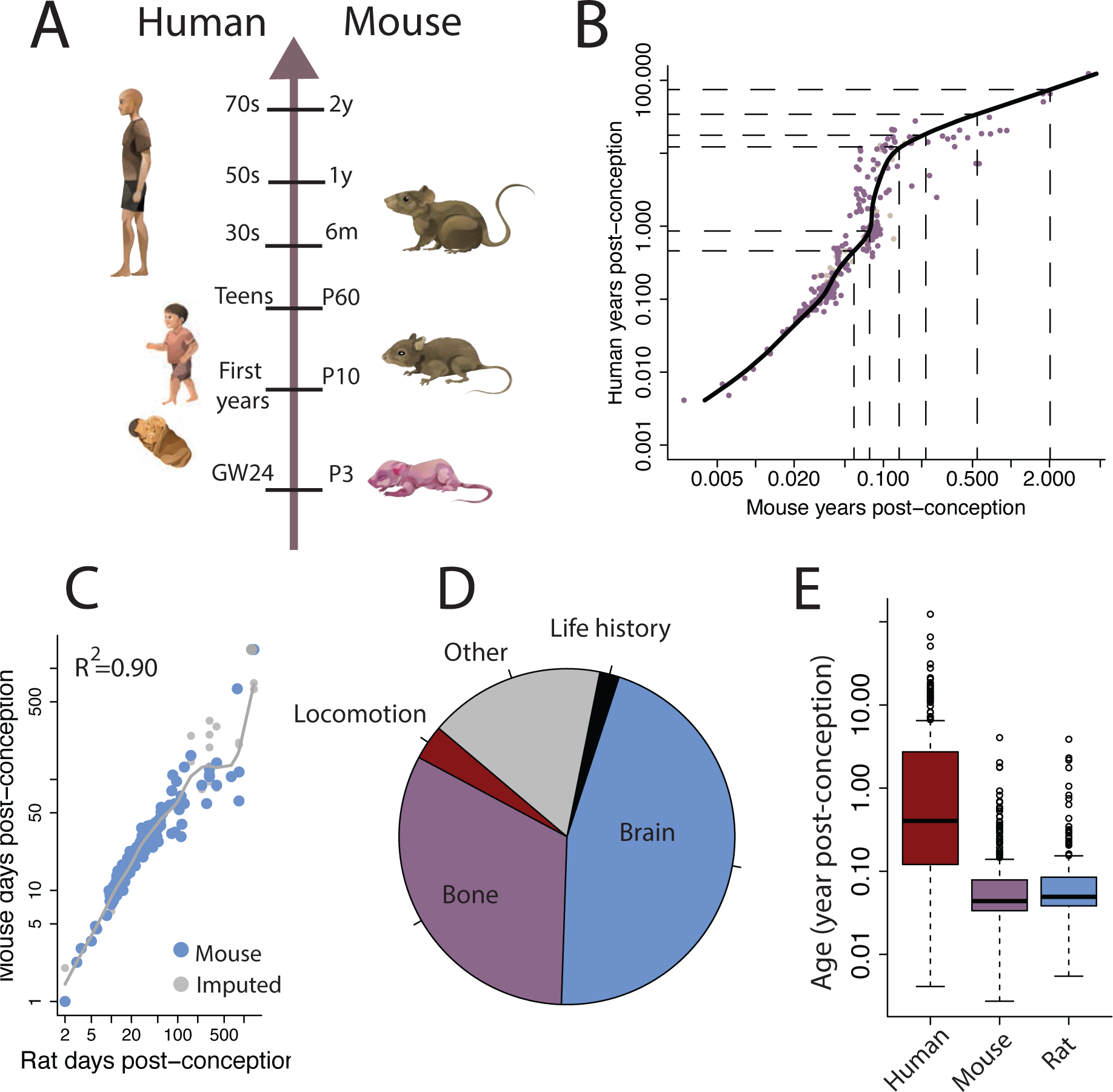
We collected time points across pre- and postnatal development in humans, rats, and mice to generate age equivalences across species. (A) According to these data, a P3 mouse equates to a human at gestational week (GW) 24, and a P10 mouse equates to a human within their first years. Moreover, a P60 mouse maps to a human in their teens, and a 2-year-old mouse maps to a human in their 70s. (B) We fit a smooth spline through time points expressed in years post-conception in humans and mice. Here, data from rats were mapped onto mice when data in mice was not available. (C) Time points in rats are plotted against those found in mice. We fit a smooth spline through these data to equate the corresponding ages of rats to mice. (D) A pie chart shows the breakdown of time point types with a majority of brain and bone time points. (E) Age ranges for these specific categories largely span the first year.

**Figure 2.**
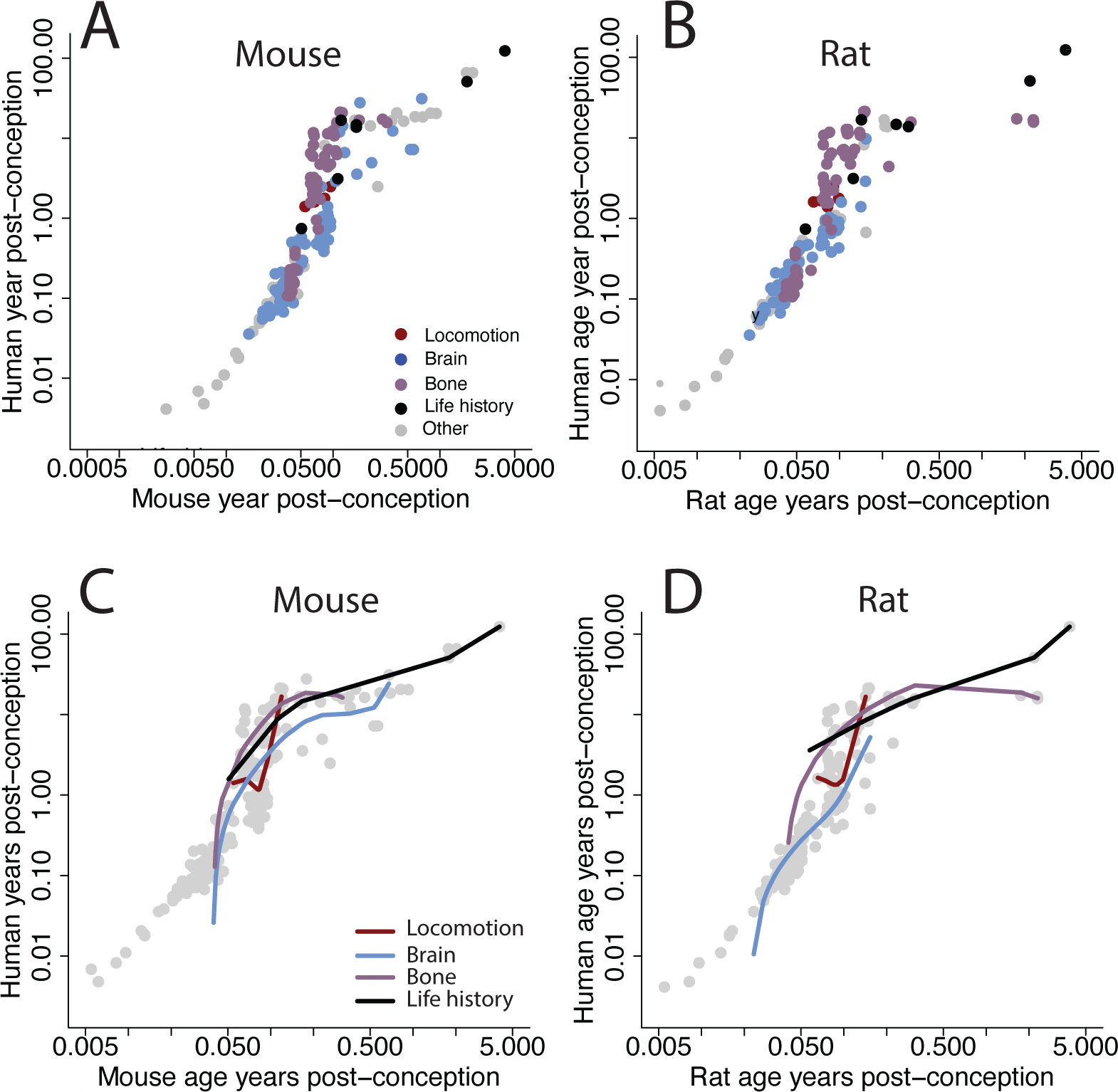
Time points (expressed in years post-conception) are plotted in (A) mice versus humans and (B) rats versus humans, which are color-coded by process. We fit a smooth spline by process to evaluate whether some processes show evidence of acceleration relative to others in (C) mice and in (D) rats. Time points show evidence of acceleration during postnatal development. All identified biological processes show relative acceleration as is evidenced by the smooth splines fit through the data.

We used a 9.4T MR scanner to collect diffusion MR scans in mouse brains. We used these data to track the postnatal growth of white matter pathways. This method relies on the diffusion of water molecules to identify the location and orientation of tracts. This information is used to reconstruct the white matter pathways of the brain in three dimensions (Mori and Zhang, 2006; Wedeen et al., 2012; Vasung et al., 2019; Charvet et al., 2020; Cottam et al., 2023). This method has been useful to characterize the development of human brain pathways across prenatal and postnatal ages. Recent work has shown that the growth of some white matter pathways largely ceases relatively soon after birth. In contrast, other pathways, such as the arcuate fasciculus, which is involved in language and higher cognition, continue to develop well into adulthood (Catani and Mesulam, 2008; Tanaka-Arakawa et al., 2015; Cohen et al., 2016; Tak et al., 2016; Mohammad and Nashaat, 2017; Vasung et al., 2019; Wilkinson et al., 2017; Wilson et al., 2021; Chen et al., 2022). We use these data to quantify the growth of association white matter pathways and we compare these growth patterns to what is known in humans. Establishing shared developmental timelines in brain circuit maturation will facilitate the translation of findings from mice to humans.

Many pathways are clearly observed in humans and mice (e.g., cingulate bundle, Schmahmann and Pandya, 2006; Wu et al., 2016), but direct comparison of white matter pathways is challenging due to substantial differences in brain structure across these two species. For example, the arcuate fasciculus is a relatively large white matter bundle that connects the frontal, parietal, and temporal lobes in humans, but there is no such clear counterpart in mouse brains (Catani and De Schotten, 2008; Catani and Mesulam, 2008; Calabrese et al., 2015; Becker et al., 2022; Charvet et al., 2022). Conversely, mice exhibit a relatively expanded olfactory association pathway traversing white matter olfactory cortical areas (see Figures 3 and 4; Collins et al., 2018), but this pathway is not distinctly observed in humans. We instead focus on comparing trajectories of multiple pathways across mice and humans. We supplement our studies of diffusion MR tractography with observations from tract-tracers, and temporal patterns in transcriptional profiles to confirm the validity of tractographies. We focus on genes expressed by long-range projections because they would provide a complimentary means to compare timelines of brain pathway maturation (Gutman et al., 2012; Jones et al., 2013; Chen et al., 2015; Johnson et al., 2019; Charvet et al., 2022; Crater et al., 2022; Cottam et al., 2023).

**Figure 3.**
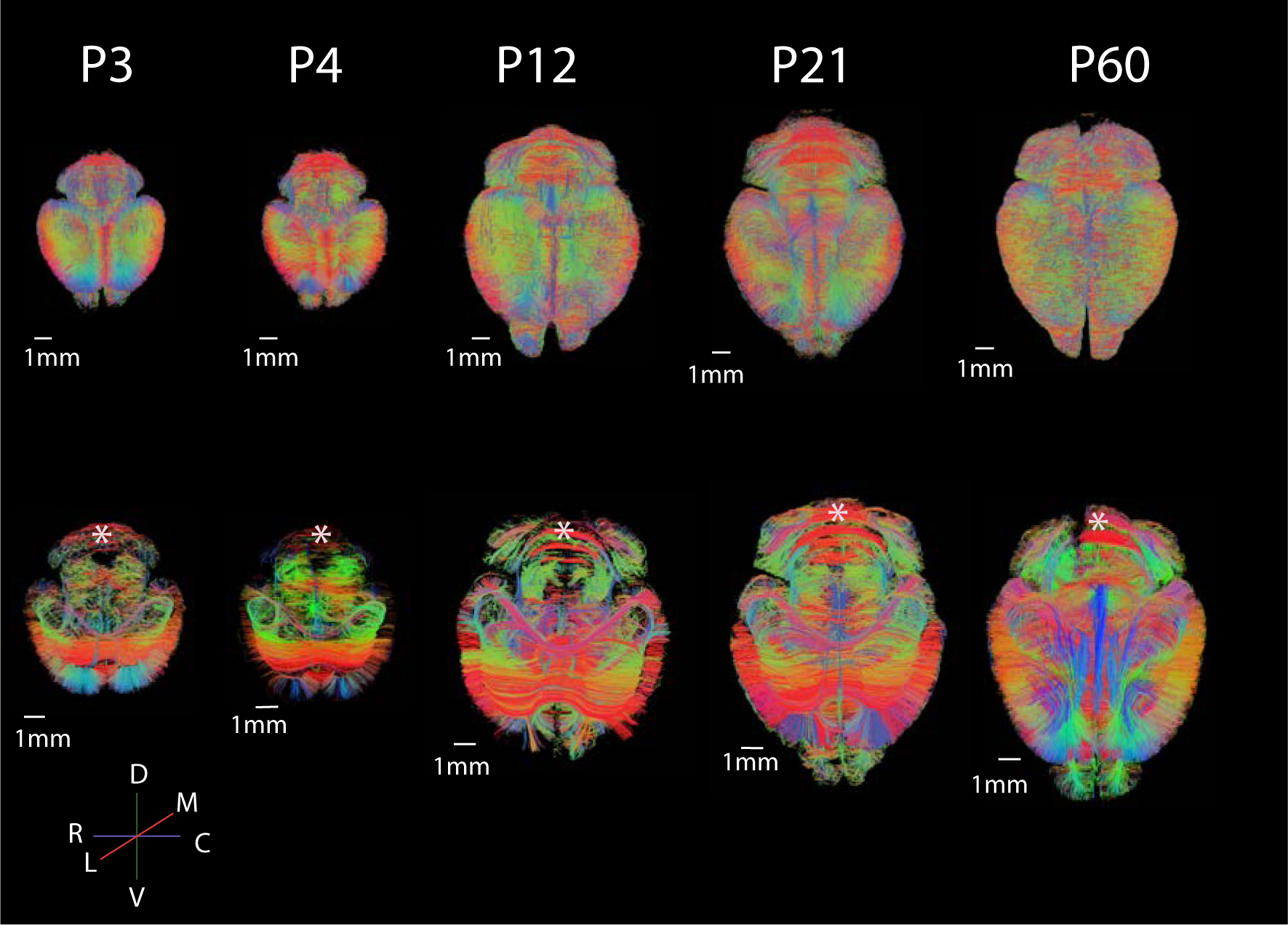
We used a 9.4T MR scanner to acquire whole-brain diffusion MR tractography. These post-mortem mouse brains were scanned at postnatal days (P) 3, 4, 12, and 60 with the color-coding representing the average direction of fibers. For example, green fibers course across the dorsal (d) to ventral (v) axis, and pathways that course across the rostral (r) to caudal (c) axis are coded in blue. In the lower panel, horizontal slices (1 slice thick) capture fibers coursing through that filter. For example, the cingulate bundle is evident, though small at birth, and expands postnatally. Cerebellar parallel fibers are evident through postnatal development (asterisk). The color-coding maps use the following abbreviations: D: dorsal; V: ventral; M: medial; L: lateral. P: postnatal day.

**Figure 4.**
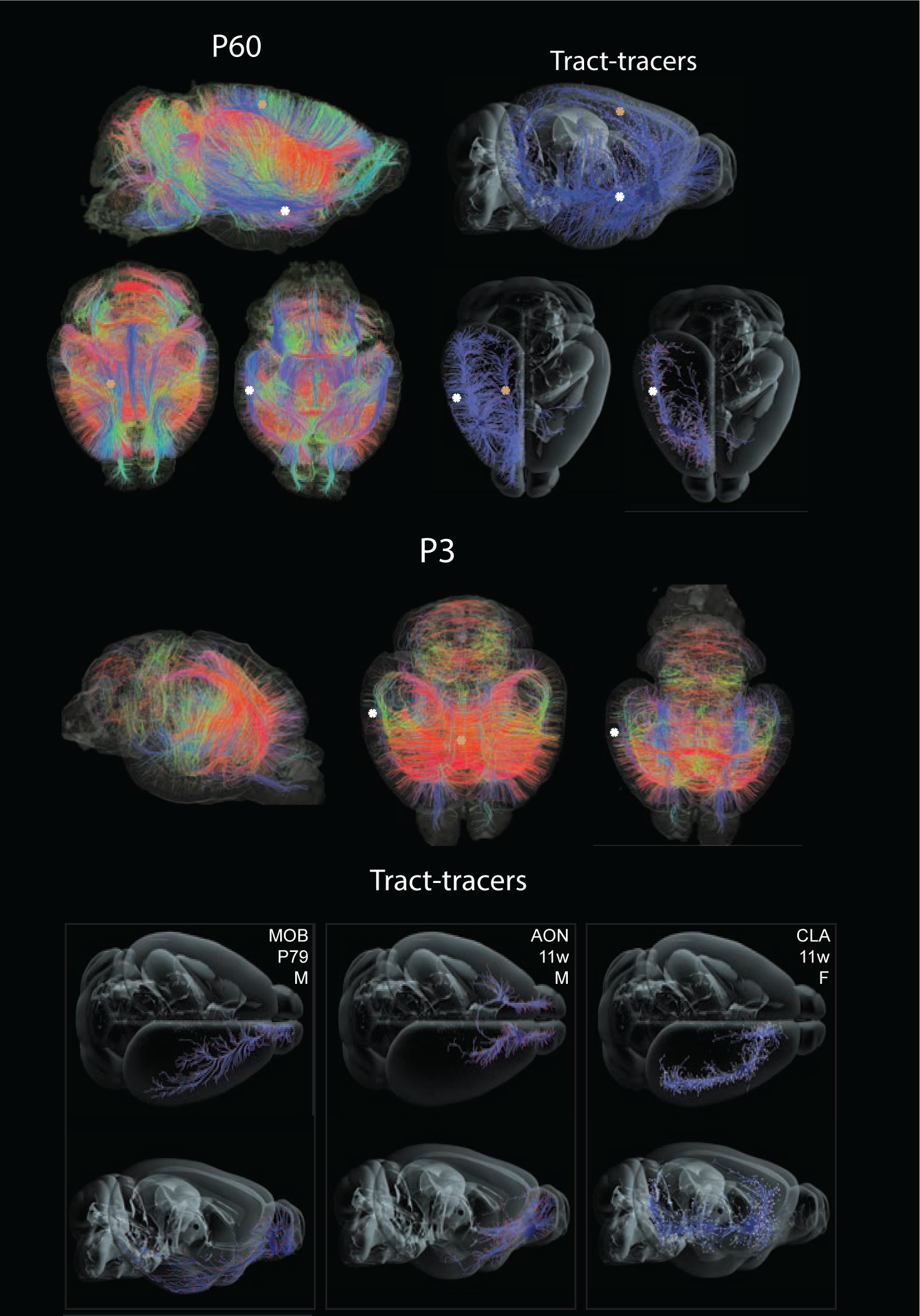
We compare diffusion MR tractography scans at P60 and P3 with tract-tracers. We found a concordance in the location and direction of fibers in the white matter across methods. Specifically, we found that P60 and P3 mouse brain diffusion MR scans show bundles of axons coursing across the rostral to caudal axis through olfactory cortices (white asterisk) and the cingulate cortex (beige asterisk). Tract-tracers label fibers that course through a similar location and direction as those found with diffusion MR tractography. Bottom panel: Additional set of tract-tracers show pathways connecting various regions, and which identify white matter pathways coursing across the rostral to caudal axis. The position and orientation of fibers resemble those observed with diffusion MR tractography. A 1-1.5 mm filter was applied to remove approximately 50% of the fibers to enable visualization of brain pathways.

## Materials and methods

### Translating time

We used a total of 477 time points (n=1,132 observations) collected across humans, rats, and mice to generate age alignments across mice and humans. We use time points collected from past studies, as well as new time points collected across pre- and postnatal ages (Figures 1-2; Workman et al., 2013; Charvet and Finlay, 2018; Charvet et al., 2017, 2023; Figure 1-1). Time points were extracted from brain, dental, bone, locomotor, and life history maturation. Examples include ages for dental eruptions, bone ossifications, sexual maturity onset, and maximum lifespan (Workman et al., 2013; Cusack et al., 2024). When maximum lifespan is omitted, mouse and human ages range from the second day of prenatal life to the third year after birth in mice, and from the second year of prenatal life to the 60s in humans (Figure 1). These time points are collected from multiple sources, which means that some variation in time points may be due to differences in procedures in data acquisition, quality, and definitions. Some variation in extrapolated ages may be due to such factors. Nevertheless, this approach is amenable to the acquisition of relatively large samples as is evidenced by a total of 1,132 observations captured across rats, mice, and humans.

Time points were extracted from abrupt and gradual changes in biological processes. We generated cross-species age alignments from temporal changes in normalized gene expression. RNA sequencing data are from the frontal cortex of humans (n=6; 35 days to 55 years old) and mice of varying ages (n=17; age range: embryonic day 11-22 months of age). These are from individuals varied in age (GSE47966; Lister et al., 2013) and were used in a past publication (Hendy et al., 2019). We selected 10 genes that could be used as markers of long-range projections (e.g., *NEFH*), synapse formation (e.g., *ARC*) as well as cell proliferation (e.g., *SOX1*) and neurogenesis markers common to both species. We captured the age at which genes peak in their expression in the frontal cortex of humans and mice. We fit non-linear regressions to age and normalized gene expression to extrapolate the age of peaks or plateaus in gene expression in both species (i.e., R library package easynls, model=2, 3). Comparing the temporal profiles in the expression of these genes provides an additional means with which to study the timeline of circuit maturation in mice as in humans.

### Translating time model

We used time points from both rats and mice to translate ages across humans and mice. Some time points are available in rats and humans, but not in mice. We used predicted age in mice from rats to increase the samples with which to equate ages across humans and mice. There were 364 time points common to mice and rats, and 54 time points available in only rats. We first imputed data with the multiple imputations by chained equations (MICE) algorithm in cases where there was one missing time point across rats and mice (Van Buuren and Groothuis-Oudshoorn, 2011). We then fit a smooth spline through the log-transformed age (expressed in days post-conception) to predict mouse time points from those available for rats (n=54; Figure 1C). The smooth spline fit accounted for 90% of the variance of these data, which means that we can with good certainty predict age in rats from mice. We used time points from rats predicted to mice when they were not available in mice.

In the present study, we focused on comparing the overall pace of development and aging across humans and mice. We do so by fitting a smooth spline to the age of mouse and human time points. This method deviates from past methods in that we do not fit linear models or quasi-Newton optimization based models (Workman et al., 2013; Charvet et al., 2023). We do not test for specific heterochronies in behavioral and biological processes across humans and mice. We know that primates and rodents show heterochronies in development. For example, it is known that cortical neurogenesis is unusually extended in primates relative to rodents (Workman et al., 2013). We consider all time points available, and we do not derive specific translations for heterochronic processes though it is possible to derive age translations for specific biological and behavioral processes (Clancy et al., 2001; Workman et al., 2013). These different approaches mean there may be some differences in specific age alignments from past approaches. We chose this approach because we are here concerned with overall age alignments rather than translating specific biological or behavioral processes.

### Specimen collection for MR imaging

Mice were maintained under a 14/10 hour light/dark photoperiod with Purina Mills Inc. (PMI) rodent diet (Animal Specialties and Provisions) and water available ad libitum. Mouse brains at postnatal days 3, 4, 12, 21, and 60 were extracted, immersion fixed in 4% paraformaldehyde, and stored at 4°C. Mouse brains were collected for purposes other than this study, which meant that this research was exempt from IACUC approval. Post-mortem mouse brains were sent to the University of Delaware Center for Biomedical and Brain Imaging. Details of the samples are in Figure 3-3.

### MR scanner

We used a 9.4 T Bruker Biospec 94/20 small animal MR system (Bruker BioSpec MRI, Ettlingen Germany) to scan post-mortem mouse brains at the University of Delaware Center for Biomedical and Brain Imaging (CBBI). Mouse brains were first immersed in phosphate-buffered saline (PBS) doped in Gadolinium diethylenetriamine pentaacetic acid (GdDTPA) for 24 hours and they were then rinsed in PBS for 6 hours then placed in fomblin before and during brain scanning. The cerebellum of some mice incurred some damage, and this damage shows in some of the scans. We acquired diffusion-weighted images using a spin-echo-based echo-planar imaging (EPI) sequence with the following parameters: an echo time (TE) of 45 ms, a repetition time (TR) of 500 ms, isotropic 100 μm^3^ voxel resolution, 65 sampling directions with a *b*-value of 4000 s/mm^2^, 5 *b* = 0 s/mm^2^ images, a slice thickness a 0.1 mm, with fat suppression and field-of-view saturation. Details on the individuals scanned, including their age, and spatial resolution are listed in Figure 3-3.

### Tractography reconstruction

We used high angular resolution diffusion imaging (HARDI) to reconstruct fiber tracts from post-mortem mouse brain diffusion weighted MR scans (Figure 3). The tractography was reconstructed with the Diffusion Toolkit (trackvis.org) to visualize and quantify pathways. Tracts were set to not exceed a 45° angle between two consecutive orientation vectors (angle threshold; Cottam et al., 2023). We used the fiber assessment by continuous tracking (FACT) algorithm to reconstruct tracks coursing through the whole brain. No fractional anisotropy threshold was applied to reconstruct tracts, which is consistent with past work (Takahashi et al., 2012; Charvet et al., 2022; Cottam et al., 2023).

### Tractography validation

Observations from tract-tracers guided the selection of tractography parameters to reduce the probability of producing inaccurate reconstructions. Indeed, it is well known that the tractography does not systematically align with findings from tract-tracers (Jones et al., 2013; Chen et al., 2015; Johnson et al., 2019; Crater et al., 2022). There are a number of reasons for such inconsistencies (Jones et al., 2013; Oh et al., 2014; Maier-Hein et al., 2017). For example, there is a likelihood that tracts mis-connect in zones of crossing fibers. In mouse brains, tracts may be misconnected at the gray to white matter cortex because of the sharp turn these fibers make at the white to gray matter boundary, though they may be less of a problem in gyrencephalic brains, which do not make such sharp turns at the white to gray matter (Charvet et al., 2022). Tractography parameters are set to filter tracts that make such sharp turns because such parameters may lead to misconnections of pathways within the white matter. These uncertainties make validation of results from tractography an important component of diffusion MR tractography studies.

Our past work showed that there is strong concordance between diffusion MR tractography and tract-tracers of commonly studied pathways. This is the case for the fornix as well as the hippocampal commissure (Charvet et al., 2022). Here, we focused on olfactory association pathways, and we used tract-tracers from the Allen Brain institute to compare tractography reconstructions with tract-tracers in the mouse brain (Figure 4). We found strong concordance in the location and orientation of white matter pathways in mice and in humans, but it was not clear whether the terminations within the gray matter are accurate. Given these observations, we measured cross-sectional areas of pathways in the white matter, and we did not focus on the precise terminations of pathways within the gray matter.

### Pathway maturation quantification

We measured the cross-sectional area of cortical and olfactory association pathways in mice from postnatal day 3 to 60 with TrackVis (Figure 3-3). We set ROIs to visualize pathways, and to measure cross-sectional pathway areas. Some fibers were measured from coronal and sagittal slices. We set ROIs through coronal slices at the level of the anterior commissure to measure the cingulate bundle and olfactory association pathway. The cingulate bundle was defined as a pathway coursing through the rostral to caudal axis of the cingulate cortex (Bubb et al., 2018). We included fibers coursing across the rostral to caudal axis of the cingulate cortex. In the medial cortex, some fibers course across the rostral to caudal and medial to lateral direction, but these were not included in the definition of the cingulate bundle. The olfactory association pathway was defined as a pathway coursing across the rostral to caudal axis of olfactory cortices. We also measured the area of the corpus callosum and the anterior commissure from mid-sagittal slices. We used these data to test whether age is a significant predictor of white matter pathway area to evaluate the timeline of growth during mouse development.

## Results

We first discuss how we equate corresponding ages across mice and humans (Figures 1-2), and how we compare the timeline of brain pathway maturation in mice relative to those of humans (Figure 4).

### Translating time across humans and mice

We collected time points (n=477) across pre- and post-natal ages in humans, mice, and rats. We extrapolated mouse time points from rat time points when data was available for both rats and humans, but was not available for mice (see methods: Figures 1-2). We fit a smooth spline through log-transformed time points (expressed in age post-conception) of mice and humans. This smooth spline fit to humans and mice accounted for 86.8% of the variance (Figure 1B), which means that we can with good certainty predict age in humans from equivalent time points in mice. We use these data to find corresponding ages across species, and to evaluate how the pace of development and aging varies with age in mice and in humans.

Humans take longer to develop than mice. The earliest time points (i.e., a 2 cell stage) occur at roughly similar ages in humans and mice (∼embryonic day 2), but later time points occur progressively much later in humans than they do in mice. For example, a mouse on postnatal day 3 equates to a human on gestation week 24, and a P10 mouse maps to a human within the first year after birth. A 1 year old mouse maps onto a human in their 50s, and a 2 year old mouse maps onto a human in their 70s. These time points find corresponding ages through much of the lifespan of humans and mice.

An interesting observation from these cross-species age alignments is that the pace at which species proceed through time points varies as a function of age. Humans take several years to proceed through milestones postnatally, but these equivalent time points only span a few months in mice. Accordingly, a P60 mouse equates to humans in their teenage years (Figure 1). We can evaluate how the pace of biological processes in humans compares with those of mice. We fit smooth splines through different biological programs to evaluate whether or multiple biological processes underlie the relative deceleration in the pace of processes in humans (Figure 2C-D). According to these smooth splines, several biological processes show relative accelerations in humans versus mice and humans versus rats (Figure 2). Those include life history, bone, and brain development, which show a relative deceleration in humans relative to mice, but such deceleration diminishes as development progresses into aging. These observations suggest the development of human infancy and childhood is relatively extended compared with mice. We next evaluate the development of pathways from diffusion MR tractography, and we use cross-species age alignments to map the timeline of white matter pathway maturation in mice compared to humans.

### Diffusion MR tractography

We collected diffusion MR scans of mouse brains (n=16) from P3 to P60 (Figure 3). Whole brain tractography shows white matter pathways, which are color-coded by the average fiber direction. At P60, many white matter pathways course across the medial to lateral axis (e.g., colossal, cortico-subcortical pathways), and others course across the rostral to caudal axis. These pathways include a relatively large bundle that spans olfactory structures named the olfactory association pathway as well as the cingulate bundle that spans the cingulate cortex. We found that the diffusion MR tractography from P3 and P60 mouse brains is concordant with tract-tracer information from mice at P56 (Figure 4). Similar to what is observed in diffusion MR tractography, tract-tracers identify axon bundles coursing across white matter olfactory structures as well as rostral to caudal cingulate fibers coursing across the cingulate cortex (Figure 4). These consistencies bolster confidence in the white matter pathway reconstructions and their analyses can be used to capture changes over the course of postnatal development.

We integrate these observations with temporal changes in gene expression (Figure 7B) and diffusion MR tractography (Figure 7 C-D). At the earliest age examined (P3), many association pathways are evident. Some pathways grow substantially during postnatal development through P60 (Figures 5 and 6). The olfactory association pathways and the cingulate bundle are relatively small at P3-4, but they expand postnatally (Figures 5 and 6). In contrast, fibers that course across the medial to lateral axis, which include the corpus callosum and the cortico-subcortical pathways, appear relatively invariant with age (Figures 5 and 6).

**Figure 5.**
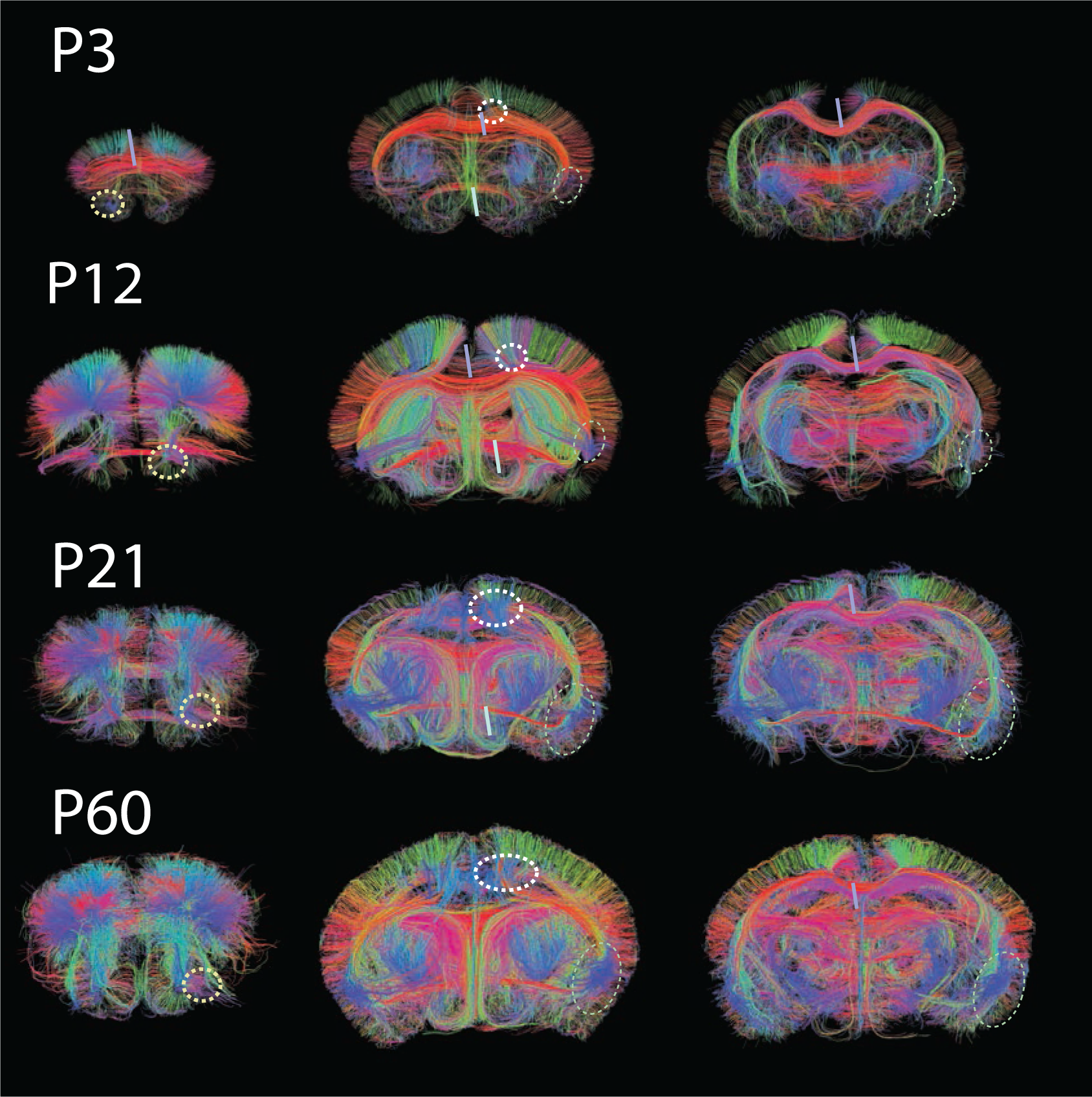
At P3, many major cortical association fibers are present. Dashed circles draw attention to the cingulate bundle as well as the olfactory association pathways, and how they grow postnatally across successive postnatal ages. A 1-slice thick coronal slice captures fibers coursing through these areas. P: postnatal day.

**Figure 6.**
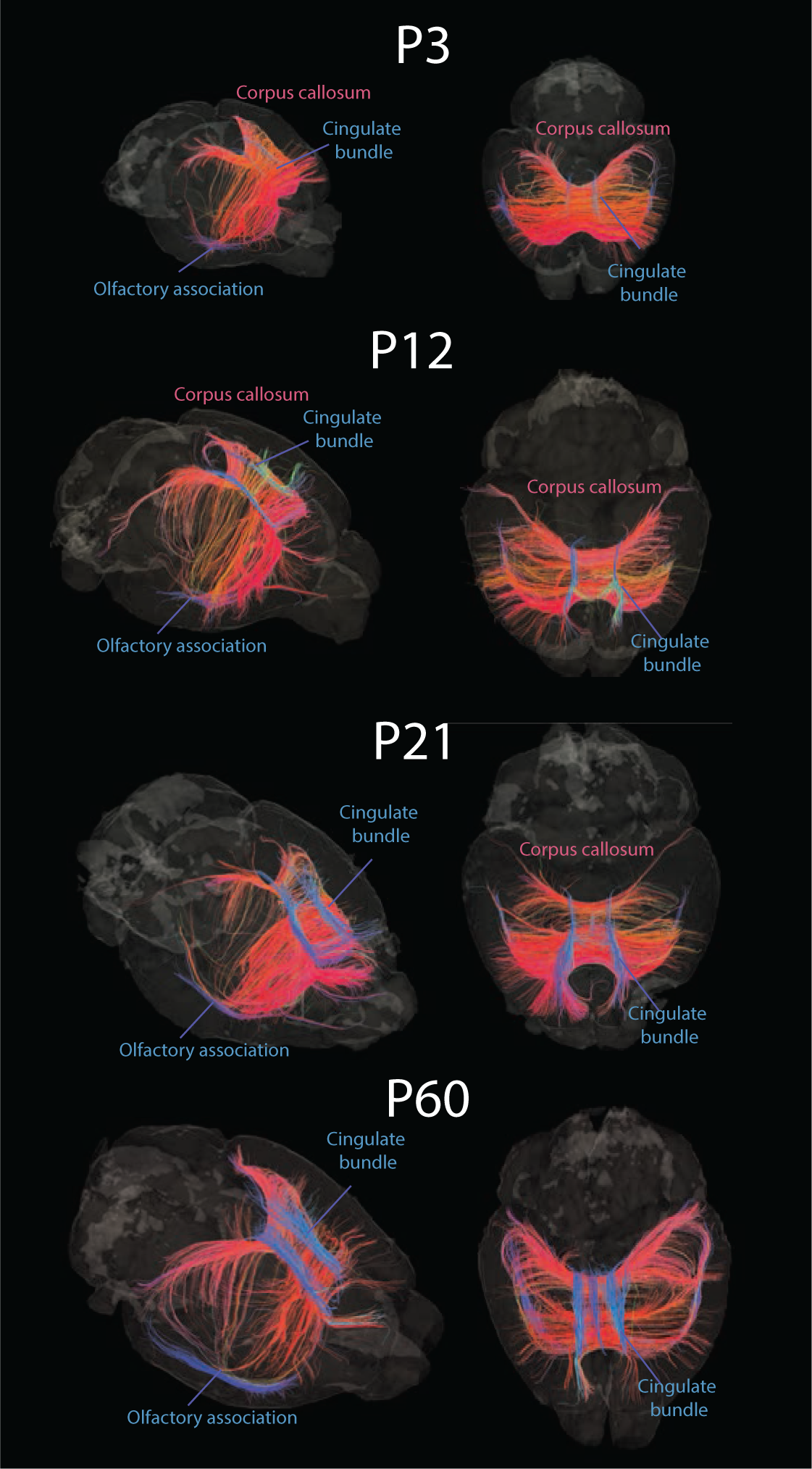
We measured the area of pathways at multiple stages of development (i.e., P3, P12, P21, and P60). We focused on measuring the area of the corpus callosum, the cingulate bundle, and the olfactory association pathway. We used ROIs to visualize coronal or sagittal slices and to quantify the area of various pathways over the course of development (Figure 7). P: postnatal days.

**Figure 7.**
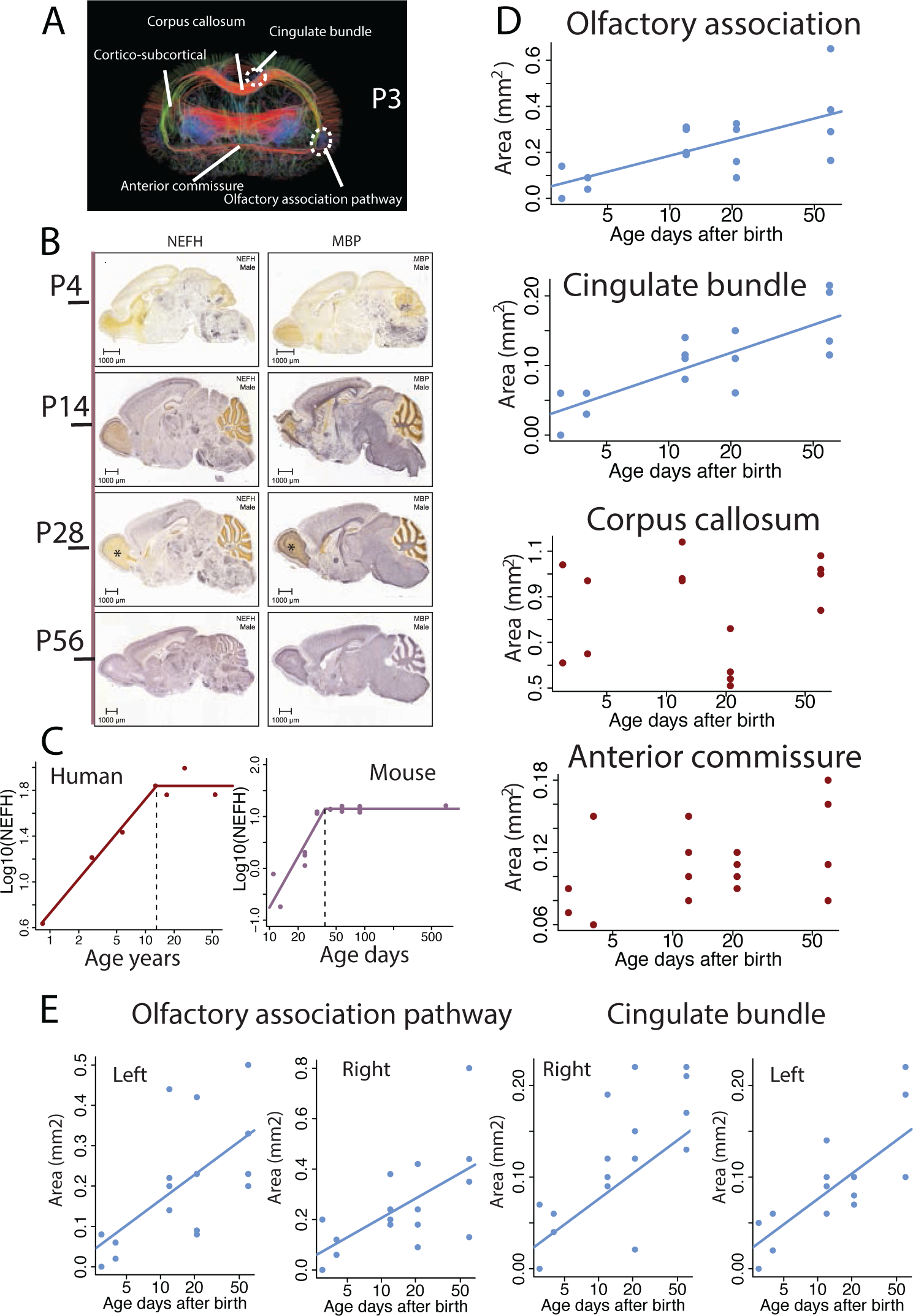
We measured the area of pathways (A), which include the cingulate bundle, the olfactory association pathways, the corpus callosum as well as the anterior commissure. (B) We also evaluated temporal changes in transcriptional profiles of *Nefh* and *Mbp* expression in mice. These are markers of long-range projecting neurons and myelination. Accordingly, *Nefh* and *Mbp* expression increases with age in mouse brains, especially between P4 and P14. The asterisk shows areas of protracted increase in expression in olfactory structures from P28 to P56. (C) We fit a nonlinear model to *NEFH* expression from RNA sequencing data from the frontal cortex of mice and humans, which show plateaus in peak expression occur at around 50 days in mice and in their teens in humans. (D) We fit linear models to test whether age could be used to predict the cross-sectional area of these pathways. (E) Left and right olfactory association pathways and cingulate bundles show sustained growth access studied ages. The olfactory association pathway and cingulate bundle increase significantly with age but this is not the case for the corpus callosum and the anterior commissure, which are relatively invariant with age. We fit linear models on these graphs when the slope of the model is significant. In situ expression data are from the Allen Brain Institute.

We fit linear models to test whether pathway area varies significantly with age (log-transformed age expressed in days post-birth; Figure 7). The olfactory association pathway (y=0.23x-0.046, n=16; adj R^2^=0.42) and the cingulate bundle (y=0.10x-0.013, n=16; adj R^2^=0.63) both significantly increase with age (slope: p=0.004 and p=0.00015, respectively). However, the anterior commissure (y=0.03x+0.07, slope p=0.11376; R^2^=0.11) and the corpus callosum (y=0.059x+0.78, slope p value=0.63; R^2^=-0.05) do not increase significantly with age. Left and right ipsilateral olfactory association pathways and cingulate bindles also show substantial growth up to P60 (Figure 7E). Together, these results demonstrate that some white matter pathways grow through P60 in mice. Given that a P60 mouse maps onto human teenage years, white matter pathways grow for an extended duration in mice as they do in humans.

### Transcription to validate pathway growth

We leveraged spatiotemporal patterns in gene expression to confirm quantitative analyses of growth trajectories made with diffusion MR tractography. We considered the spatiotemporal expression of Myelin Basic Protein (*MBP*) and Neurofilament Heavy Polypeptide (*NEFH*) with in situ expression because these genes are expressed by large neurons with long axonal structure and can be used as markers of neurons with long-range projections (Zečević et al., 1998; Khalil et al., 2018). There is a sharp increase in the expression of *MBP* and *NEFH* from P4 to P14 (Figure 7B). The expression of *MBP* and *NEFH* increases in some structures up to at least P56 and this includes olfactory structures (Figure 7B). Such increases are also observed from RNA sequencing data, which show peak expression in *NEFH* in the frontal cortex in the first postnatal month (P19.5) and teens in humans (12.3 years post-birth). A similar situation is observed with frontal cortex *NEFH* expression, which increases up to around 50 days in mice and in their teens in humans (Figure 7C). These data further confirm that white matter pathways mature well into human teenage years, which equates to the second postnatal month in mice.

## Discussion

In this study, we leveraged Translating Time as well as neuroimaging techniques to assess spatiotemporal changes in the growth of murine white matter pathways for comparison with humans. We also generate cross-species age alignments in humans and mice, which can serve as a resource for researchers who need to map findings across species (Clancy et al., 2007). Additionally, this study demonstrates that diffusion MR tractography is a valuable tool to characterize the brain’s white matter tracts in model systems, especially once the tractography is integrated with other methods to validate the tractography.

### Translating time

We expand on our long-term research project called Translating Time, which equates corresponding ages across model systems and humans (Clancy et al., 2001; Workman et al., 2013; Charvet and Finlay, 2018; Charvet et al., 2023). We incorporate time points from multiple metrics to find corresponding ages during pre- and postnatal development. In humans, biological processes that span some postnatal ages, including neural circuit maturation and bone ossification, proceed more slowly at postnatal ages than they do during prenatal stages. This relatively lengthened pace of postnatal development supports the notion that humans have a relatively extended duration of childhood and helplessness during which infants can absorb vast amounts of information. Learning without the need for immediate task performance results in a powerful foundation model that supports rapid and versatile learning later on (Cusack et al., 2024). These time translations enrich our understanding of conserved as well as modified biological processes leading to phenotypic variation.

There is much interest in identifying equivalent timelines across mice and humans. Such age translations are important to study human age-related diseases that emerge at late stages of life, including Alzheimer’s disease (Westergaard et al., 2019). It’s an open question as to how old mice should be to model human aging. Mice do not show the full spectrum of age-related diseases found in humans (Marx et al., 2013; Rigby Dames et al., 2023). For example, mice do not spontaneously develop plaques and tangles, which is a hallmark of Alzheimer’s disease phenotypes nor do they typically develop cataracts, and mice rarely live past 18 months of age (Fahlström and Ulfhake B, 2011). Since mice do not spontaneously develop many of the symptoms for human aging, they are genetically modified to recapitulate some aspects of these diseases. Our cross-species age alignments illuminate which mouse age is most useful for studying age-related diseases in humans. The alignments we’ve developed extend to 70 years of age in humans, which equates to 2 years of age in mice. Given that Alzheimer’s disease risk substantially increases in the 70s and beyond, our study suggests that a comparable age in mice extends beyond 2 years of age though very few mice live sufficiently long lives to map onto human geriatric stages.

### Diffusion MR tractography captures white matter pathway growth

Diffusion MR can be used to reconstruct brain pathways, and should only be paired with other biological approaches such as gene expression, histology, tract-tracers, or other techniques (Axer et al., 2011; Gutman et al., 2012; Calabrese et al., 2015; Aydogan et al., 2018; Winnubst et al., 2019; Axer and Amunts, 2022; Charvet, 2023; Charvet et al., 2022; Johnson et al., 2023). Here, a comparison of tract-tracers and tractography reconstructions was used to guide the tractography reconstruction parameters (Chen et al., 2015; Martin-Lopez et al., 2018; Charvet et al., 2022; Cottam et al., 2023). White matter association pathways grow for an extended duration in mice. These data show that brain pathways mature for an extended time at postnatal ages in both mice and humans.

There are substantial differences in wiring across species. For example, humans possess several cortical association pathways (e.g., arcuate fasciculus, inferior longitudinal fasciculus), which are linked to higher cognitive capacities and language. Many of these cortical association pathways are found in apes and monkeys, but they become progressively more elusive in increasingly distant species. Indeed, many of the association pathways found in humans are also elusive in mice. Mice also possess association pathways (e.g., olfactory association pathways) that do not have a clear homology to humans. Given these observations, we did not focus on comparing specific pathways across humans and mice. Instead, we focus on overall trajectories in white matter pathway development in mice, and how such timelines compare to humans.

We found heterogeneity in white matter pathway maturation in mice, which is a situation akin to what is found in humans (Cohen et al., 2016; Mohammad et al., 2017). In mice, the corpus callosum and the anterior commissure do not grow significantly at postnatal ages whereas the olfactory association pathway and the cingulate bundle grow well through P60. Our age alignments across humans and mice show that P60 maps onto human teenage years. Therefore, our study shows that some white matter pathways extend to similar timelines across both humans and mice.

While the overall timelines are similar across species, our study also highlights interesting cross-species differences. Here, mice show protracted development of limbic structures, including the cingulate bundle and olfactory association pathways. In humans, cortical association pathways, such as the arcuate fasciculus, mature for a relatively extended duration. In humans, these pathways are linked to higher cognitive processing and some of our species-specific adaptations, including language, are especially protracted throughout development, and their growth extends well into our twenties. The protracted development of limbic pathways in mice is linked to murine specializations for olfaction. These observations raise the intriguing possibility that protracted development of pathways may be a signature of sensory or cognitive species-specific specializations.

## Conclusions

Translating Time is a powerful tool that provides an easily accessible method to correlate studies between animal models and humans. We collected 477 time points related to brain, dental, bone, and locomotor maturations, as well as lifespans from previous studies and we equated them across humans, rats, and mice. We also use diffusion MR tractography to identify key tract characteristics in white matter pathways such as the cingulate bundle and corpus callosum in P3, P12, P21, and P60 mice, and map the developmental timeline to their equivalencies in humans. The growth of the Translating Time tool is providing researchers with important translational context to increase the impact the utility of model systems, and the reliability of cross-species experimentation and comparison.

## Data availability

Diffusion MR scans and cross-species time points will be made available with the publication of this work. The scripts and the data will be available on publication. The P60 and P21 mouse diffusion brain scans used in this study are available on Dryad.

## Acknowledgments

This work was supported by Eunice Kennedy Shriver National Institute of Child Health and Human Development Grants 1R21-HD-101964-01A1 and 7R21-HD101964-02 (to C.J.C.); Institutional Development Award Networks of Biomedical Research Excellence Pilot Grant P20-GM-103446 from the National Institute of General Medical Sciences (NIGMS; to C.J.C.); NIGMS Core Center Access Award P20-GM-103446 (to C.J.C.); startup funds from Auburn University (to C.J.C.), and an Auburn undergraduate research fellowship (to M.B). The Centers of Biomedical Research Excellence Grant 5P20-GM-103653 was used for research at Delaware State University; Opinions are not necessarily those of the National Institutes of Health.

## Author contributions

Study concept: CJC, JS; Data collection: KO; NC, JS, KH; Data analysis: MB, CJC, NC, JR; Statistical analyses: CJC; Writing: CJC; Editing: KO; NC, JS, KH, JR; Funding: CJC.

